# Advancing spatially explicit fisheries management with age-specific species distribution models

**DOI:** 10.1101/2025.02.27.640560

**Authors:** Eric J. Ward, Nick Tolimieri

## Abstract

Many species distribution models (SDMs) incorporate external information, such as environmental or habitat features, yet the majority overlook age-specific information. Including age information may be particularly valuable for species exhibiting age-related spatial patterns driven by ontogenetic shifts, recruitment dynamics, and selective fishing pressures. Here, we develop an age-structured SDM framework using data from the west coast of the USA and concentrate on two species with distinct life histories: North Pacific hake (*Merluccius productus*) and sablefish (*Anoplopoma fimbria*). We validate our approach by forecasting age classes across survey locations in future years; these results highlight that forecast models have predictive skill (sablefish more so than hake) and that the predictive ability is highest for older age classes. Using predictions of age-1 sablefish as an example, we demonstrate how our models may be used to forecast future bycatch risk in space, helping increase the efficiency of fisheries. Our framework is applicable to a wide range of survey types and platforms around the world and supports the development of spatially explicit management strategies and optimal allocation of fishing effort.

## Introduction

Species distribution models (SDMs) are widely used in marine ecology to understand and predict the spatial distribution of species based on environmental conditions, mechanisms, and biological processes (Melo-Merino et al. 2020). These models have informed management decisions for conservation and resource allocation, especially in the face of changing environmental conditions and human impacts, such as fishing pressure (Elith and Leathwick 2009; Guisan and Thuiller 2005). Recent advances in SDM applications in the marine environment have incorporated external drivers such as temperature (Fredston et al. 2021) and habitat features (Laman et al. 2018; Phillips et al. 2017). A second use of these models has been to predict temporal trends in species biomass from spatiotemporal fisheries independent survey data (Thorson et al. 2015). Trends estimated from SDMs may be included in integrated population models (stock assessments) for commercially fished species or used to identify risks to populations of conservation concern. Recent developments in these models have included depth (Johnson et al. 2019) and habitat variables (Cao et al. 2017) to improve trend estimation.

While many SDMs around the world have included variables such as temperature, depth, or habitat, they have generally overlooked age structure, and instead modeled either species occurrences or densities across all age or size classes. Omitting information on age or size may be partially dependent on data availability (ageing fish generally requires sampling otoliths, which is costly, and surveys may not observe younger age classes for some species), but may also be driven by the modeling choices of analysts. Much of the previous research has demonstrated that age structure is a fundamental characteristic of fish populations, influencing spatial distribution and abundance. Mechanisms may include ontogenetic shifts (including shifts in depth or space; Dahlgren and Eggleston 2000), recruitment variability (particularly for species whose recruitment is strongly linked to the environment), and selective fishing pressures (Rijnsdorp et al. 2018). Fishing pressure may alter the age composition of a population, and because there is a spatial component of fishing effort, these impacts may be spatial (Ono et al. 2016). Recent work has also highlighted that age-based SDMs can improve understanding of environmental drivers on commercially valuable populations (Ono et al. 2024). Ignoring age-specific information in SDMs may lead to incomplete or inaccurate predictions, especially for species with strong age-related spatial gradients.

Incorporating age structure into SDMs may provide more accurate predictions by capturing spatial and temporal variability related to life history traits, age-specific habitat preferences, and population dynamics (Karp et al. 2025). Fish often exhibit ontogenetic shifts, where younger individuals occupy different habitats than adults due to factors like predator avoidance, habitat preferences, and diet changes (Ciannelli et al. 2004; Werner and Gilliam 1984). These shifts can create age-specific distributions across habitats, particularly for species that undergo distinct life history phases. Additionally, spatiallly variable recruitment pulses and age-related migration patterns can influence the spatial distribution of younger versus older fish. Fishing selectivity often further accentuates these patterns by disproportionately removing larger, older fish, which alters the population structure and potentially shifts spatial distributions (Hilborn and Walters 1992; Methot and Wetzel 2013). By incorporating age, SDMs could better predict these dynamics and improve our understanding of species-environment relationships.

Understanding the spatial distributions of age classes also is important for both conservation and fisheries management (Kopf et al. 2024). Previous work has linked population productivity to age structure, with above average contributions from larger and older females (Ohlberger et al. 2022). For populations where spatial distribution varies by age or size, identifying spatial concentrations of larger, older females, is useful for identifying areas associated with elevated productivity (Berkeley et al. 2004). Understanding how the spatial age structure of a population intersects with the spatial allocation of fishing effort and fishers’ targeting behavior also has implications for fisheries management. Though results vary by fisheries and gear, for some fisheries larger sized individuals are more valuable per kg than small or medium sized individuals (Tsikliras & Polymeros 2014).

Because of the recursive nature of population dynamics, e.g. numbers at age *a* at time *t* can be used to predict numbers at age *a*+1 at time *t*+1, age-structured SDMs with predictive skill also provide a tool for forward prediction, with considerable implications for marine resource management. Reliable predictions of species distributions can help managers anticipate and mitigate issues such as bycatch risk by identifying regions with high densities of specific age cohorts (Lewison et al. 2004). Forecasting the spatial distribution of certain age classes can inform spatially explicit management strategies and help optimize fishing effort to target sustainable cohorts. For example, knowing the expected future locations of specific age classes could aid in creating time-area closures that protect vulnerable life stages or ensure harvests focus on more resilient portions of the population. However, building robust age-structured SDMs requires high-quality, long-term datasets with consistent sampling methods across both age and space to ensure the validity of these predictions.

Around the world, age and life history information is routinely collected by a number of sampling platforms, including from fisheries and fisheries independent surveys. While fisheries dependent data are often used to inform age structured models, these data may include biases due to preferential spatial sampling or gear selectivity; data from fisheries independent surveys may also include size- or age-selectivity, but these effects are more consistent across space and time. The primary objective of our paper is to explore the utility of developing age - structured SDMs from fisheries independent data. Focusing on two species with contrasting life history characteristics in the North Pacific, the semi-pelagic North Pacific hake (*Merluccius productus*) and longer – lived benthic sablefish (*Anoplopoma fimbria*), we first develop a flexible framework for modeling spatially explicit numbers at age. These species also represent contrasts in ontogeny; older sablefish generally move offshore to deeper depths (Head et al. 2024), while Pacific hake tend to migrate north as they age (Agostini et al. 2006). Second, we evaluate the ability of this framework to offer predictive skill in future forecasts and quantify how the predictive relationship changes as a function of age. Finally, we demonstrate how model output may be used to examine changing species compositions through time, make maps of potential bycatch hotspots, and identify areas with higher concentrations of mature fish as potential targets of fishing effort.

## Methods

### Data

On the west coast of the USA, the West Coast Groundfish Bottom Trawl Survey (WCGBTS) has been an annual survey conduced from 2003 - present. The WCGBTS is designed to estimate the abundance, size, and age composition of groundfish species important to commercial and recreational fisheries found in near-bottom habitats on the west coast of the USA (Keller et al. 2017). The survey effort is concentrated in summer months and has been conducted annually since 2003 (data are publicly available at https://www.nwfsc.noaa.gov/data). Importantly, the random stratified sampling design, spatial and seasonal coverage, effort, and gears have remained constant within the period we analyze. Like many other surveys around the world, no WCGBTS survey occurred in 2020 because of the Covid-19 pandemic. Though the WCGBTS survey samples hundreds of species, we concentrated our analysis on the two species with the most age data from the WCGBTS, North Pacific hake and sablefish. These species are also commercially important, and previous work indicates different ecologies related to the distribution of juvenile fishes with hake having more spatially variable juvenile distribution from year to year (Tolimieri et al 2020). While otoliths are sampled continuously during the WCGBTS survey for a broad range of species, otoliths are generally aged when species are being prioritized for stock assessment by the Pacific Fishery Management Council (PFMC); full assessments between species may occur irregularly and be sporadically updated every 5 – 10 years. Hake and sablefish represent exceptions, as the hake stock is assessed annually by co-managers from the USA and Canada (Grandin et al. 2024) and sablefish is frequently assessed because of its high commercial value. For our analysis, we used sablefish collected 2003 – 2023 (except for 2020 when there was no survey; mean 1314.4 individuals sampled per year; Table S1) and Pacific hake collected 2007 – 2019 (mean 650.7 individuals sampled per year; age information was not collected for 2004-2006 or from 2020 – present; Table S2). We filtered the data to include only female fishes because typically more females are aged than males and because stock assessments estimate female spawning biomass. We also truncated ages to focus on those with the highest data availability; for hake this included ages 1 – 5, and for sablefish ages 0 – 9 (data from these ages represented 54% and 75% of the total aged fish, respectively).

### Statistical modeling

We constructed a unique spatiotemporal model for each species – age combination in our dataset. We adopted this approach partially because of smaller sample sizes relative to applications that have modeled ages simultaneously (Ono et al. 2024). An advantage of modeling all ages simultaneously is that it allows for additional complexity (such as shared year effects between age classes). However an advantage of modeling ages independently is greater flexibility (e.g. unique autocorrelation and spatial variance parameters by age). Individual fish were aggregated at the haul level. As these aggregated counts are zero-inflated (85% of hake and 89% of sablefish observations) and represent small counts (< 7), we modeled counts at age with a Binomial distribution. This approach also incorporates variability in sampling effort, using the total number of fish sampled as the Binomial N parameter. We constructed a spatiotemporal model as an extension of Generalized Linear Mixed Models (GLMMs) such that the prediction in location *s* and time *t* can be written as

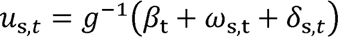

where **u*_s,t_* represents the mean predicted probability of occurrence, *g*^-l^() represents the inverse link function (logit), *β_t_* represent time – varying intercepts modeled as an AR(1) process 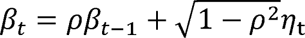, where 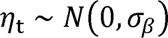, the spatial field **ω** ∼ Multivariate Normal(0, Σ_ω_) and the spatiotemporal fields ***δ**_t_* are modeled as an AR(1) process 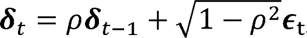, where ***∈*_t_** ∼ MVNormal(**0, Σ**_*∈*_). The spatial field **ω** represents the average spatial distribution across years, while the spatiotemporal field ***δ_t_*** quantifies interannual variation in the spatial distribution likely in response to environmental variables (Thorson 2019), and in our parameterization, includes temporal autocorrelation (the AR(1) process).

Spatial and spatiotemporal random fields were constructed as Gaussian Markov random fields (GMRFs) using the Stochastic Partial Differential Equation approach (SPDE) (Lindgren et al. 2011; Lindgren and Rue 2015). The SPDE method models the spatial correlation between points as a Matérn covariance function with smoothness parameter *v = 1*. Spatial meshes for all ages and species were constructed with a cutoff distance of 50km (the cutoff distance controls the spacing of mesh vertices). Parameter estimation was done using the sdmTMB software package (Anderson et al. 2024) with R 4.3.1 (R Core Team 2024). The sdmTMB package relies on Template Model Builder (TMB; Kristensen et al. 2016) to quickly and efficiently maximize the marginal log likelihood using auto-differentiation and the Laplace approximation to integrate out random effects. Models were evaluated for convergence (positive-definite Hessian matrix, and a maximum absolute log likelihood gradient < 0.001) and residuals diagnostics were evaluated with the DHARMa package (Hartig, 2022). Code and data to replicate our analysis is publicly available on Github, https://github.com/noaa-nwfsc/spatial-age-models.

### Validating future predictive ability

As a first validation, we leveraged the natural recursive element of our data to quantify the ability of our models to predict the future distribution by age. For each of the species - age models constructed above, we made predictions to the survey locations in the following year (e.g. age 3 hake in years 2007 – 2018 was used to predict the distribution of age 4 hake in 2008 – 2019). For each year, we related the predicted probabilities of occurrence 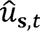 to observations by multiplying predictions by the total number of fish sampled for ageing (**N*_s,t_*) in the validation dataset, 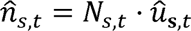. We quantified the relationship between predictions and observations by fitting a simple Poisson GLM, where counts of fish of age *a* in year *t* were treated as the response and 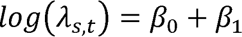 · 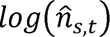. The exponentiated slope parameter **exp(β**_1_)represents a change in expected counts that would be expected from a 1 – unit change in 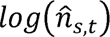. As a diagnostic check and to understand whether there were potential temporal patterns in the response, we fit a second model as a GLMM with R package glmmTMB (Brooks et al. 2017); this second model was identical to our Poisson GLM with random effects in *β*_0_ and *β*_l_, where year was used as the grouping variable.

### Using predictions to quantify bycatch risk

To illustrate potential benefits to fisheries management, we used output from our models to generate spatiotemporal predictions of age – 1 sablefish in year *t* (from age – 0 sablefish in year *t* - 1). Sablefish often occur in an assemblage including Dover sole *Microstomus pacificus*, shortspine and longspine thornyheads (*Sebastomus alascanus and S. altivelis*, respectively). This complex (known as ‘DTS’) forms an important fishery on the West Coast, and bycatch of sablefish can impact quotas for the other three species, as well as the at–sea hake fishery. For example, in 2017, the at–sea hake fishery had 50 mt of sablefish quota set aside in the area north of 36°N, but the fleet caught more than 3 times as much, prompting in-season warnings to alter fishing behavior.

Gridded predictions from our Poisson GLM represent the expected number of fish per unit of effort in each location. To turn these into a potentially more useful measure of risk, we calculate the predicted total age – 1 sablefish catch per unit of effort (CPUE) within a given radius of major fishing ports on the US west coast. For consistency with previous work, we used a radius of 282 km (Leising et al. 2024). These predictions of port-level CPUE were then summarized as annual time series (1 – step ahead forecasts).

## Results

One useful output from our spatiotemporal models is the estimated spatial field, *ω_s_*, representing the average spatial distribution across years. Results from our model of North Pacific hake show a concentration of age – 1 hake near the coast, and more of latitudinal north – south break for 2 – 3 year old hake (with higher occurrences in the South, Fig. 1). As expected, age 4 hake appear to have a northern distribution, though the gradient is less clear than for ages 2 – 3. For sablefish, our models estimate a concentration of age 0 – 2 sablefish in shallow water, with a deeper distribution by age 3 (Fig. 2). Latitudinally, there are several areas in central Oregon that appear to have consistently lower occurrences than average for older age classes (Fig. 2).

**Figure 1:**
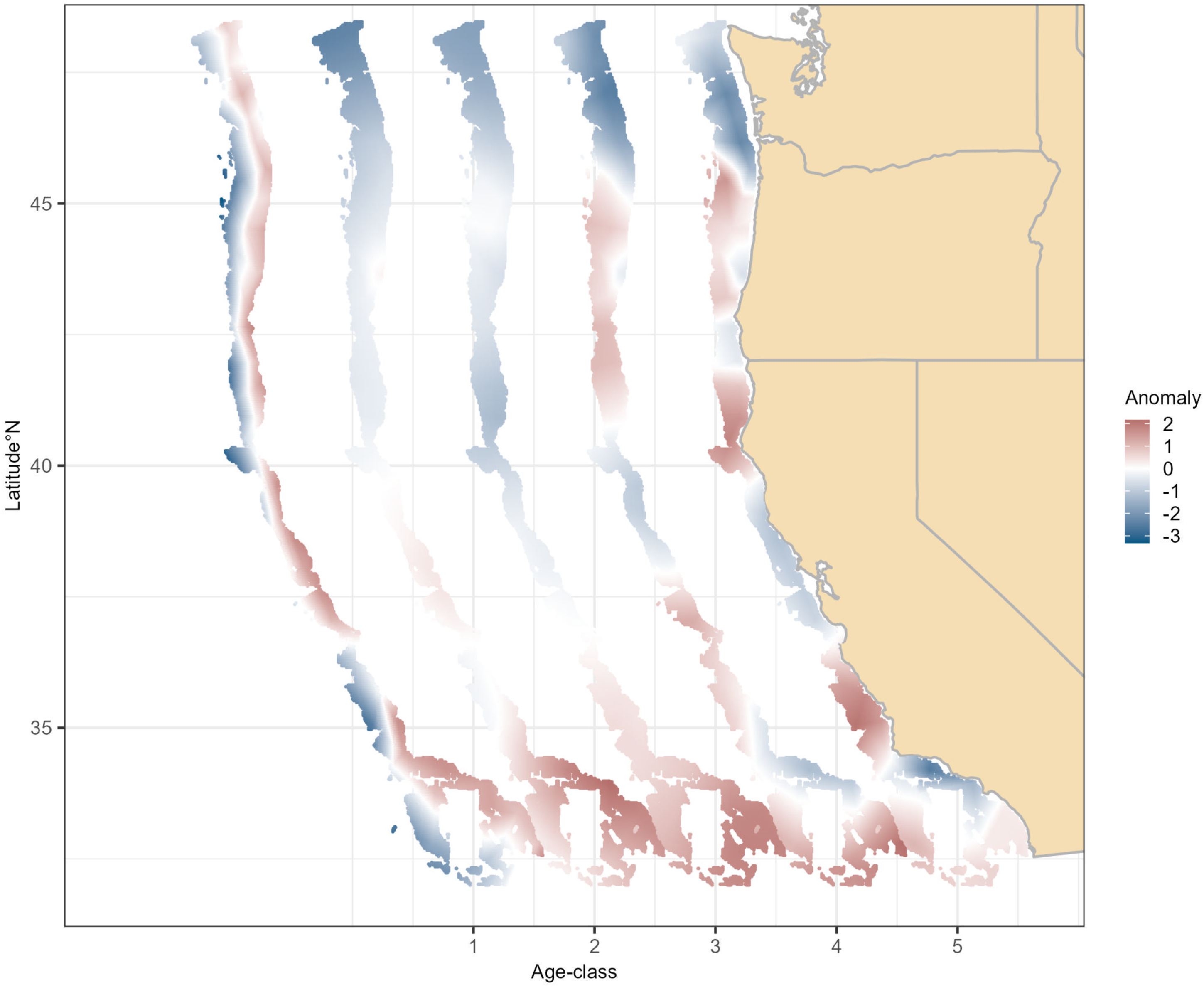
Estimated spatial anomalies for Pacific hake ages 1 – 5. These fields represent the average anomalies across all years, 2003 – 2023. The spatial anomalies for each age class have been shifted along the longitude (x) axis to present the age-specific anomalies on the same figure.

**Figure 2:**
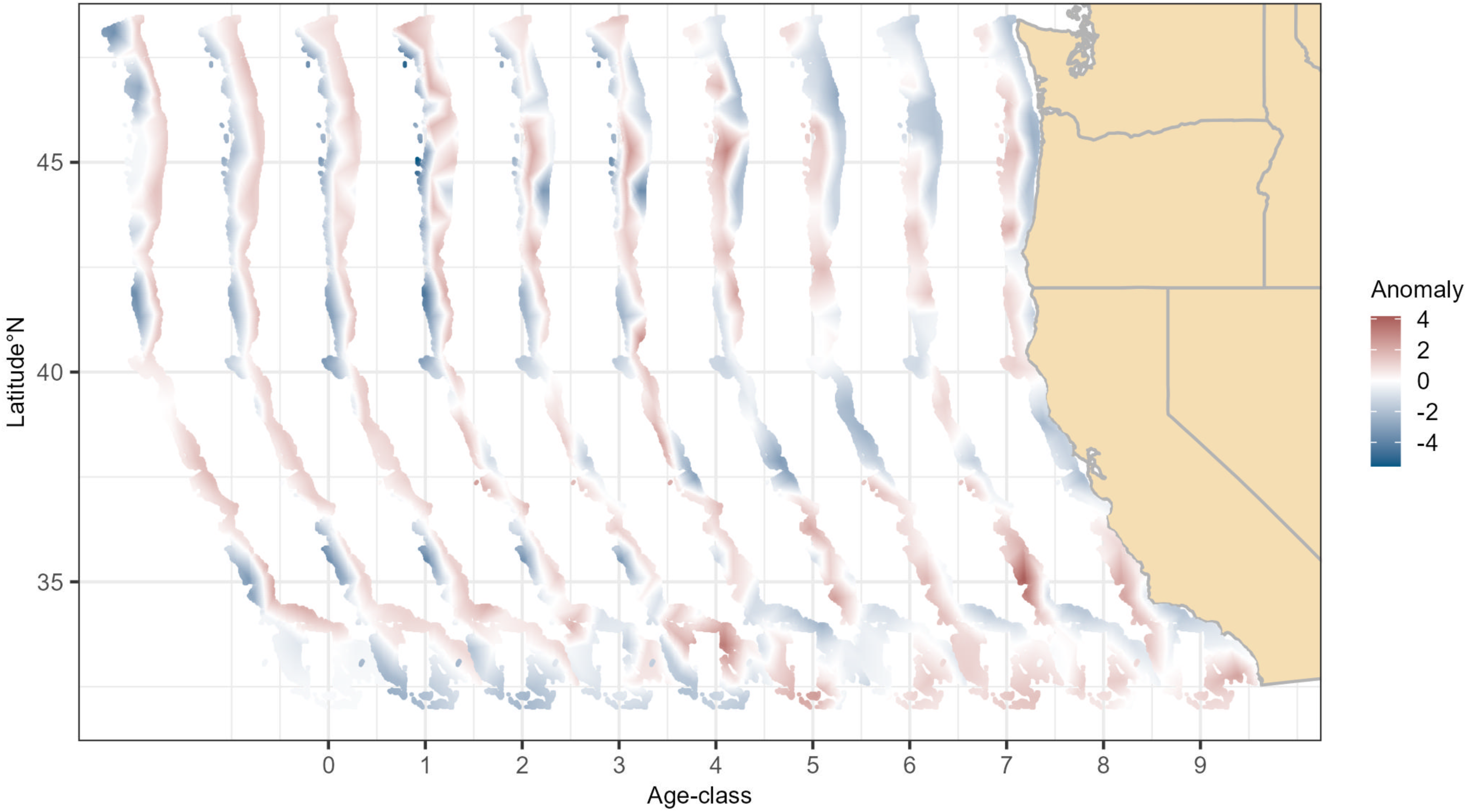
Estimated spatial anomalies for sablefish ages 0 – 9. These fields represent the average anomalies across all years, 2003 – 2023. The spatial anomalies for each age class have been shifted along the longitude (x) axis to present the age-specific anomalies on the same figure.

### Tracking strong age cohorts

Estimates of age composition by year may be useful in tracking the distribution of such cohorts through time, as well as examining variability in distributions of particular age classes. For example, for both hake (Figs. 3 and S2) and sablefish (Figs. 4 and S3), distinct patches of high age – 0 abundance tended to disperse as the fish aged such that spatial distributions for older age–classes were less distinct. While predictions can be made for any cohort, strong cohorts represent both future fishing opportunities and potential bycatch issues. Therefore, we also used the results from our models to project the spatial distributions of strong hake cohorts (2008, 2010, 2014, 2016; Fig. 3 and Fig. S2) and strong sablefish cohorts (2008, 2010, 2016, 2021, Fig. 4 and Fig. S3). These predictions are also useful in revealing subtle differences between strong cohort years. For example, the 2014 cohort of Pacific hake had a larger fraction of age – 1 individuals distributed in the northern part of the domain (along the coast of Washington state), while other cohorts had hotspots in the center and southern portion of the domain. Similarly for sablefish, the 2016 cohort was concentrated in the north (age–1 individuals in 2011), versus other cohorts that had a more coastwide distribution. In fact, age–1 fish were uncommon south of approximately Cape Mendocino (40°N) from 2015 – 2020 (Fig. S3), concurrent with marine heatwaves during this period.

**Figure 3:**
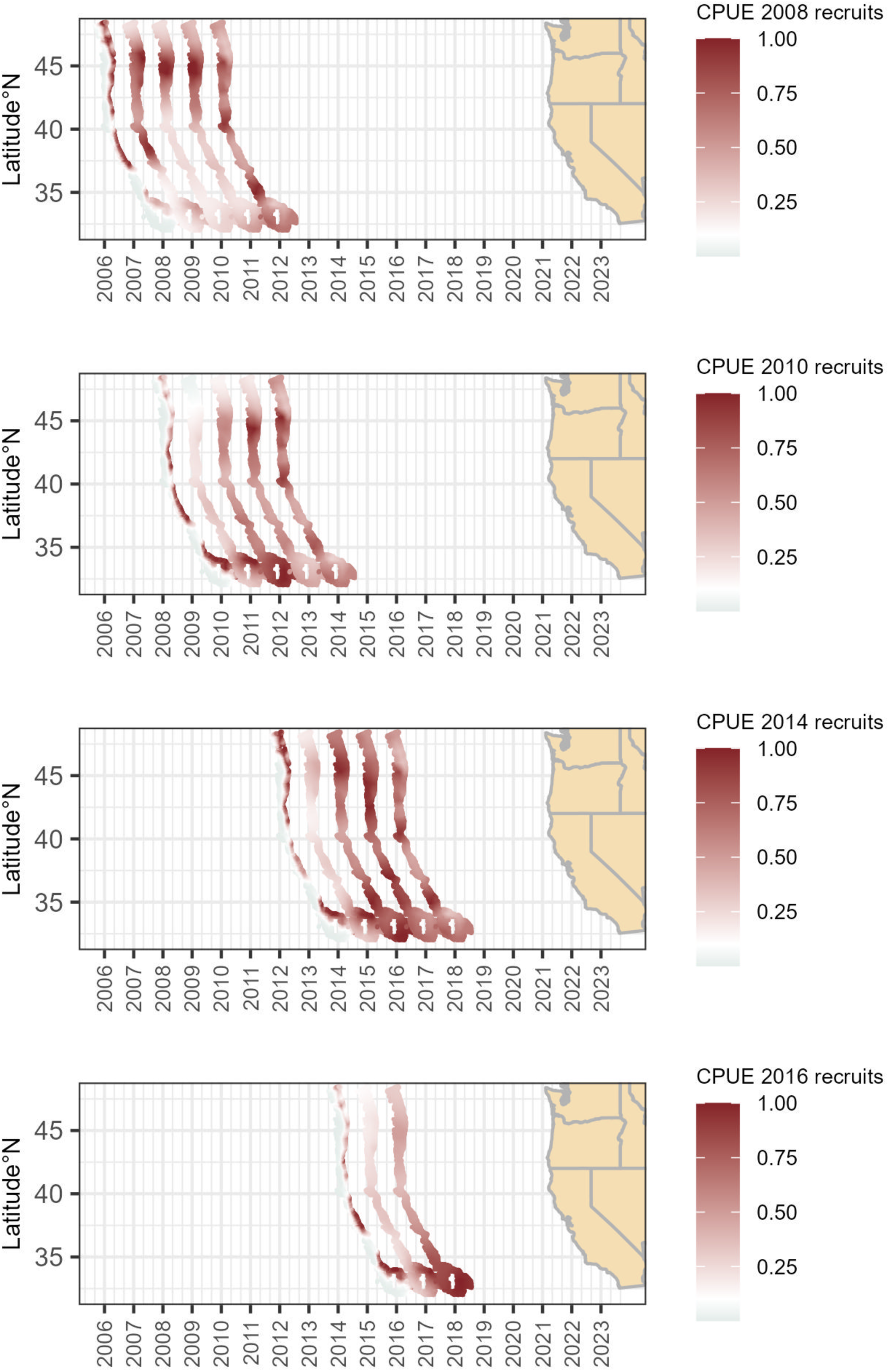
Estimated spatial catch per unit effort (CPUE, kg per km^2^) for Pacific hake; rows represent strong cohorts in our dataset (2008, 2010, 2014, 2016). CPUE is standardized to 1.0 across cohorts for visualization purposes. Full predictions for all cohorts are in the Supplementary Information. The CPUE maps for each year have been shifted along the longitude (x) axis to present the age-specific anomalies on the same figure.

**Figure 4:**
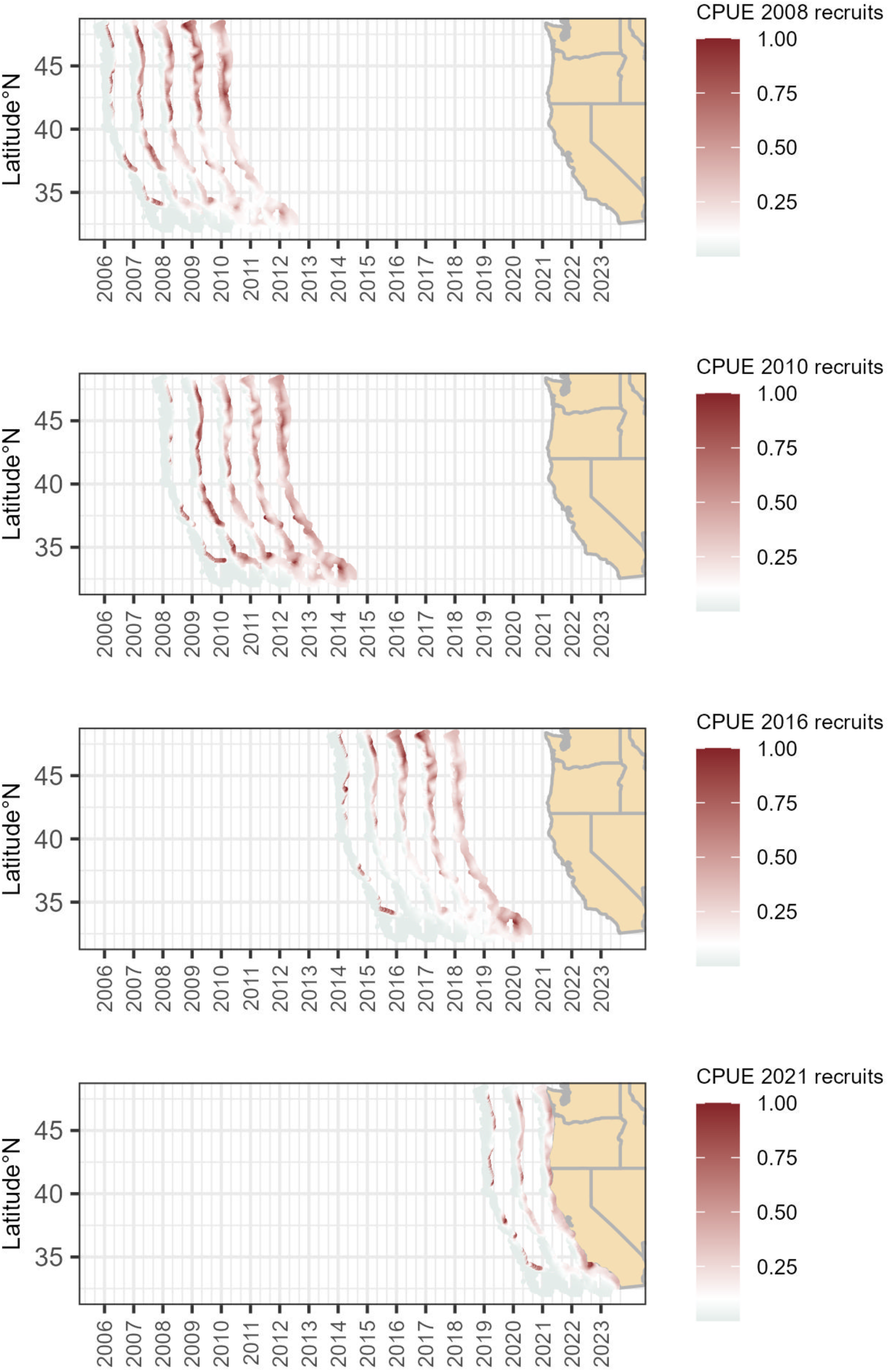
Estimated spatial catch per unit effort (CPUE, kg per km^2^) for sablefish; each row represents a strong cohort in our dataset (2008, 2010, 2016, 2021). CPUE is standardized to 1.0 across cohorts for visualization purposes. Full predictions for all cohorts are in the Supplementary Information. The CPUE maps for each year have been shifted along the longitude (x) axis to present the age-specific anomalies on the same figure.

When comparing the spatial and spatiotemporal parameters across ages and species, we find that for most ages, there is a decrease in the estimated spatial and spatiotemporal variances as well as a decrease in the spatial range with increasing age (Fig. S1). Spatial and spatiotemporal variances control the magnitude of the peaks and valleys in estimated latent fields, meaning that spatial fields estimated for older fish are less variable and more spatially homogenous. Similarly, the spatial range determines the distance at which locations are functionally independent (for the Matern, *ρ* ∼ 0.13). The decline in range with age implies that spatial distributions also become more patchy as a function of age (Fig. S1).

### Forecasts

Our 1–step ahead validation analyses indicate that there is a stronger association between older ages compared to younger ages, and this effect is strongest for sablefish (Fig. 5). When comparing our initial Poisson GLM to a more complicated GLMM with random year effects in the intercept and slope (Fig. S4), we found little support for shared correlations across age classes of a given species. Shared correlations would indicate for example that age – 2 and age – 3 sablefish may be more or less predictable in certain years (indicating that information about one age class may be useful in predicting another). With the exception of a strong positive correlation between age – 0 and age – 1 sablefish, most correlations between age classes were smaller than 0.3 in magnitude (Fig. S4).

**Figure 5:**
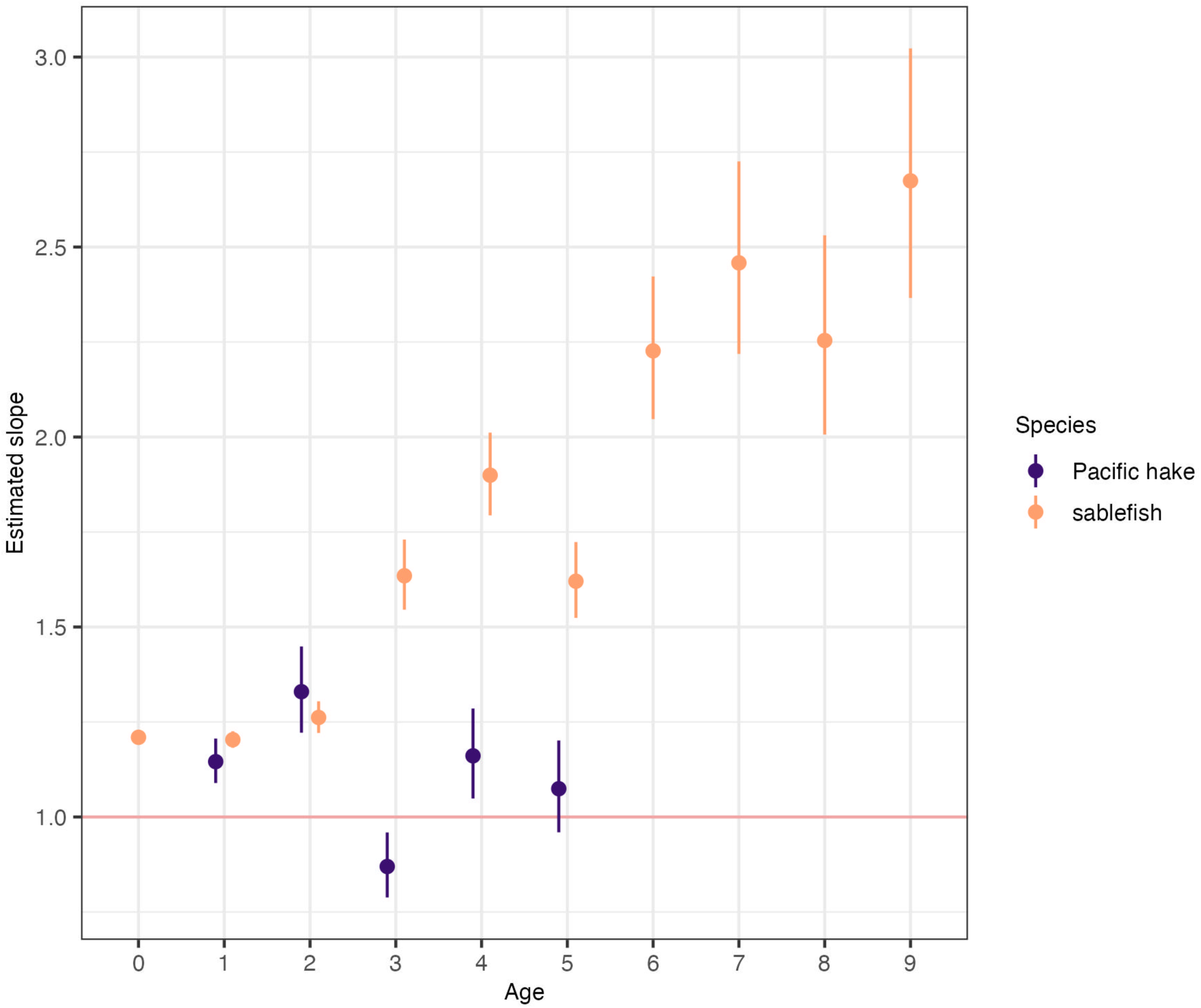
Estimated coefficients relating predicted densities of age a fish in year t to observed numbers the following year. The coefficients and 95% confidence intervals (vertical lines) are estimated in log space and presented here in normal space.

### Quantifying bycatch risk of juvenile sablefish

Our forecasts of age – 1 sablefish CPUE illustrates the variability of potential bycatch risk in space and time. In most years for example, age – 1 CPUE is lowest in ports in the state of California (Eureka, Fort Bragg, Morro Bay) and highest in several ports in the state of Oregon (Astoria, Coos Bay, Fig. 6). These results also illustrate the high temporal variability in age – 1 CPUE, with peaks occurring in 2009, 2014, 2017, and 2021, Figure 6). During peak years, there is also variability among ports – forecasts of age – 1 CPUE were similar for Astoria and Coos Bay in 2021 for instance, but the potential bycatch risk near Astoria was higher in 2017 (Fig. 6).

**Figure 6:**
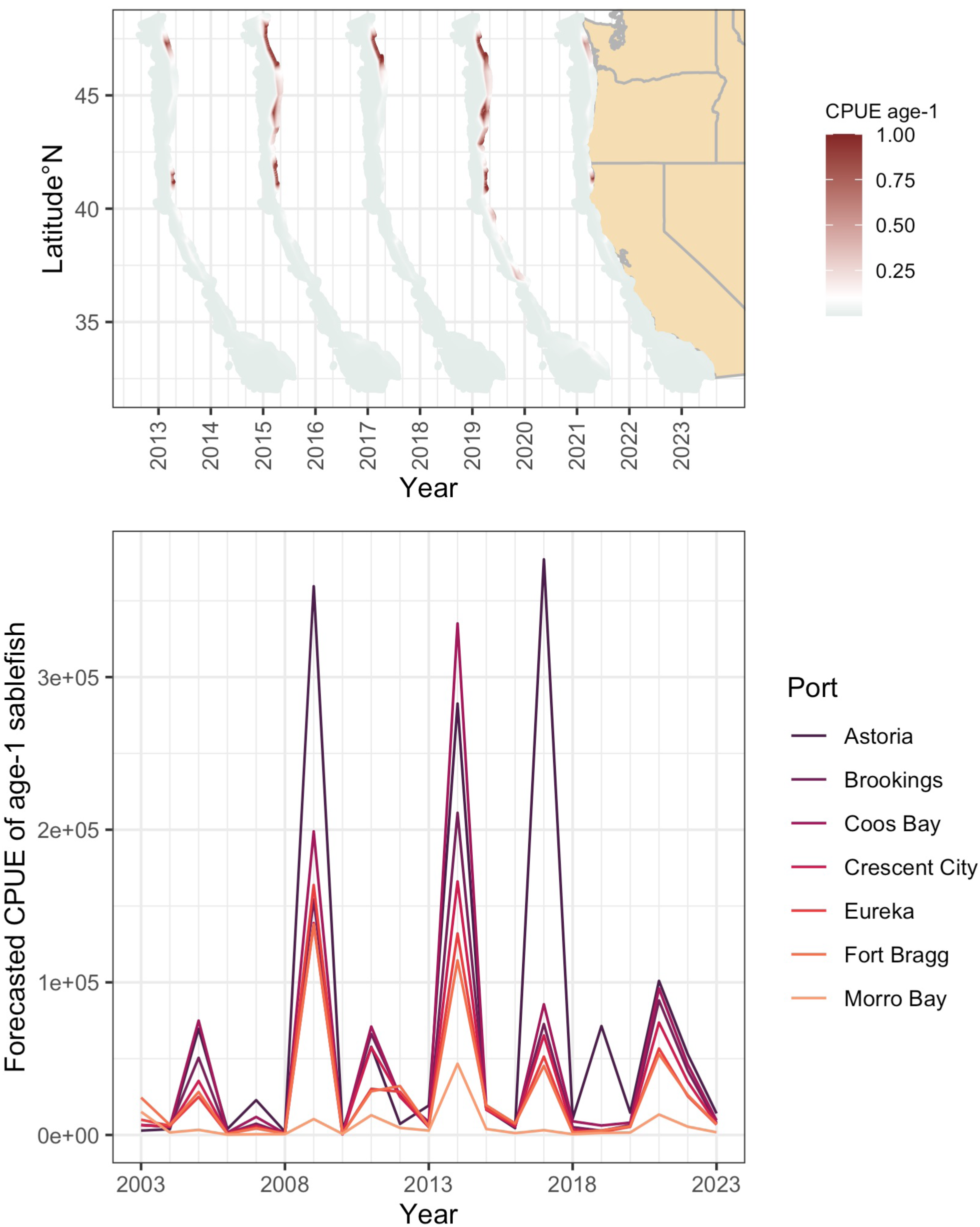
Estimated spatial catch per unit effort (CPUE, kg per km^2^) for age-1 sablefish and port-level forecasts of age-1 CPUE within a radius of 232 km. In the upper plot CPUE is standardized to 1.0 across cohorts for visualization purposes, and maps for each year have been shifted along the longitude (x) axis to present the age-specific anomalies on the same figure.

## Discussion

In this study, we developed an age–structured species distribution modeling (SDM) framework to explore spatial age composition and predict age–specific distributions of two commercially important fish species with different growth rates and spatial distributions: North Pacific hake and sablefish. Our approach integrates spatiotemporal dynamics, age structure, and forward–predictive capabilities, providing a useful tool for fisheries management. By examining age–specific patterns in spatial distributions and validating predictive skill, we highlight the potential utility of incorporating age structure into SDMs for advancing spatially explicit fisheries management.

### Age-Structured Spatial Patterns

Results from our age–structured spatial models underscore the importance of accounting for age in SDMs. The spatial patterns observed for both hake and sablefish were consistent with their distinct life histories, supporting the idea that ontogenetic shifts significantly influence fish distributions. Younger age classes of hake and sablefish exhibited broader distributions generally in shallower areas, while older individuals were more concentrated offshore (sablefish in deeper waters, hake in more northern regions). The deeper distribution of older sablefish reflects their preference for benthic habitats and an ontogenetic shift to deeper waters, while the observed north-south gradient in older hake aligns with known migratory behaviors influenced by spawning and feeding grounds (Agostini et al. 2006).

### Predictive skill and cohort tracking

Our validation analyses revealed that the predictive skill of our framework increases with age, particularly for sablefish. This result is intuitive given that older age classes generally exhibit more stable spatial patterns due to reduced recruitment variability and more predictable habitat preferences (Head et al. 2014). In contrast, the lower predictive skill for younger age classes reflects their higher spatiotemporal variability, likely driven by recruitment pulses and early–life survival dynamics. Tracking cohorts through time using our framework proved effective in identifying strong year classes (Figs. 3 – 4). Identifying these cohorts may be useful to stock assessment and management – for instance, understanding the spatial distribution of strong cohorts could be used to inform spatial harvest allocation (Olsen et al. 2012). Additionally, the ability to project spatial age composition provides a tool for assessing how environmental changes, such as ocean warming or habitat shifts, may impact the distribution of key cohorts over time.

### Managing bycatch risk

Our case study using age–1 sablefish demonstrates the practical utility of our modeling for mitigating bycatch risk. Forecasts of CPUE for age–1 sablefish revealed significant temporal and spatial variability, with higher bycatch risk near Oregon ports in certain years. Importantly, these forecasts could be provided to industry partners and fisheries managers 6 – 12 months before fishing occurs. Such information could help fishers avoid areas associated with high bycatch risk or be used to adjust the spatial distribution of quota in certain years. The observed peaks in age–1 sablefish CPUE during certain years suggest the influence of recruitment pulses, which are thought to be tied to environmental drivers (Tolimieri et al. 2018; Tolimieri and Haltuch 2023). Future efforts could also incorporate environmental covariates to enhance its predictive accuracy and further support climate–resilient fisheries management.

### Need for reliable age data

Perhaps the biggest challenge in implementing our modeling framework in the future is that these models depend on reliable age data being collected. While the WCGBTS provides a robust dataset, gaps in sampling (e.g., the absence of a 2020 survey due to the COVID–19 pandemic) and variability in otolith aging efforts over time could introduce uncertainty. The costs of ship-based surveys, such as the WCGBTS survey, have increased around the world, and numerous efforts are being implemented to make these programs more efficient (ICES 2020). While age samples are more expensive to collect than length information, there is value from the increased precision associated with age data. For data limited situations, it may be beneficial to aggregate age classes (or examine correlations between ages), or explore the use of stage structured models.

### Future directions

Our age–structured SDM framework represents a significant advance in spatial modeling, offering novel insights into the dynamics of fish populations and their age-specific distributions. From a management perspective, age–structured SDMs offer a powerful tool for addressing critical challenges, including bycatch mitigation, spatial planning, and sustainable harvesting. By providing high–resolution predictions of spatial age composition, these models enable managers to align fishing effort with ecological and economic goals, ensuring the long–term sustainability of marine resources. Future research could extend this framework by incorporating environmental covariates, such as temperature or oxygen, to account for dynamic habitat preferences, or fisheries removals to account for local depletion (Ono et al. 2016). Additionally, exploring the potential for multi–species models could provide insights into ecosystem-level interactions and their influence on spatial distributions.

## Acknowledgments

We thank Owen Liu, Chris Jordan, and 2 anonymous reviewers for constructive comments on the manuscript.

## Author contributions

EJW and NT contributed equally to all aspects of this manuscript, including conceptualization, formal analysis, visualization, and writing.

## Data availability statement

All data and for replicating this analysis are publicly available on our Github repository, https://github.com/noaa-nwfsc/spatial-age-models, and on publication will be archived at Zenodo.

## References

Agostini, V. N., Francis, R. C., Hollowed, A. B., Pierce, S. D., Wilson, C., Hendrix, A. N. 2006. The relationship between Pacific hake (*Merluccius productus*) distribution and poleward subsurface flow in the California Current System. Canadian Journal of Fisheries and Aquatic Sciences, 63(12), 2648–2659. 10.1139/f06-139

Anderson, S. C., Ward, E. J., English, P. A., Barnett, L. A. K., Thorson, J. T. 2024. sdmTMB: An R package for fast, flexible, and user-friendly generalized linear mixed effects models with spatial and spatiotemporal random fields. bioRxiv. 10.1101/2022.03.24.485545

Berkeley, S. A., Hixon, M. A., Larson, R. J., Love, M. S. 2004. Fisheries sustainability via protection of age structure and spatial distribution of fish populations. Fisheries, 29(8), 23–32. 10.1577/1548-8446(2004)29[23:FSVPOA]2.0.CO;2

Brooks, E., M., Kristensen, K., Benthem, A. van, K. J. Magnusson, Berg, C. W., Nielsen, A., Skaug, H. J., Mächler, M., Bolker, M., B. 2017. glmmTMB balances speed and flexibility among packages for zero-inflated generalized linear mixed modeling. The R Journal, 9(2), 378. 10.32614/rj-2017-066

Cao, J., Thorson, J. T., Richards, R. A., Chen, Y. 2017. Spatiotemporal index standardization improves the stock assessment of northern shrimp in the Gulf of Maine. Canadian Journal of Fisheries and Aquatic Sciences, 74(11), 1781–1793. 10.1139/cjfas-2016-0137

Ciannelli, L., Chan, K.-S., Bailey, K. M., Stenseth, N. Chr. 2004. Nonadditive effects of the environment on the survival of a large marine fish population. Ecology, 85(12), 3418–3427. 10.1890/03-0755

Dahlgren, C.P., Eggleston, D.B. 2000. Ecological processes underlying ontogenetic habitat shifts in a coral reef fish. Ecology, 81: 2227–2240. 10.1890/0012-9658(2000)081[2227:EPUOHS]2.0.CO;2

Elith, J., Leathwick, J. R. 2009. Species distribution models: Ecological explanation and prediction across space and time. Annual Review of Ecology, Evolution, and Systematics, 40(1), 677–697. 10.1146/annurev.ecolsys.110308.120159

Fredston, A., Pinsky, M., Selden, R. L., Szuwalski, C., Thorson, J. T., Gaines, S. D., Halpern, B. S. 2021. Range edges of North American marine species are tracking temperature over decades. Global Change Biology, 27(13), 3145–3156. 10.1111/gcb.15614

Grandin, C. J., Johnson, K. F., Edwards, A. M., Berger, A. M. 2024. Status of the Pacific Hake (whiting) stock in U.S. and Canadian waters in 2024. Prepared by the Joint Technical Committee of the U.S. and Canada Pacific Hake/Whiting Agreement, National Marine Fisheries Service and Fisheries and Oceans Canada. 246 p.

Guisan, A., Thuiller, W. 2005. Predicting species distribution: offering more than simple habitat models. Ecology Letters, 8(9), 993–1009. 10.1111/j.1461-0248.2005.00792.x

Hartig, F. 2022. *DHARMa: Residual diagnostics for hierarchical (multi-level / mixed) regression models*. https://CRAN.R-project.org/package=DHARMa

Head, M. A., Keller, A. A., Bradburn, M. 2014. Maturity and growth of sablefish, Anoplopoma fimbria, along the U.S. West Coast. Fisheries Research, 159, 56–67.

Hilborn, R., Walters, C. J. 1992. Quantitative fisheries stock assessment. Springer US. 10.1007/978-1-4615-3598-0

ICES. 2020. Workshop on unavoidable survey effort reduction (WKUSER). ICES. 10.17895/ICES.PUB.7453

Johnson, K. F., Thorson, J. T., Punt, A. E. 2019. Investigating the value of including depth during spatiotemporal index standardization. Fisheries Research, 216, 126–137. 10.1016/j.fishres.2019.04.004

Karp, M.A., Cimino, M., Craig, J.K., Crear, D.P., Haak, C., Hazen, E.L., Kaplan, I., Kobayashi, D.R., Moustahfid, H., Muhling, B., Pinsky, M.L., Smith, L.A., Thorson, J.T., Woodworth-Jefcoats, P.A. 2025. Applications of species distribution modeling and future needs to support marine resource management, ICES Journal of Marine Science, 82(3), fsaf024, 10.1093/icesjms/fsaf024

Keller, A. A., Wallace, J. R., Methot, R. D. 2017. *The Northwest Fisheries Science Center’s West coast groundfish bottom trawl survey: History design, and description*. NOAA, U.S. Department of Commerce. 10.7289/V5/TM-NWFSC-136

Kopf, R.K. et al. 2024. Loss of Earth’s old, wise, and large animals. Science387, eado2705. DOI:10.1126/science.ado2705

Kristensen, K., Nielsen, A., Berg, C. W., Skaug, H., Bell, B. M. 2016. TMB: Automatic differentiation and Laplace approximation. Journal of Statistical Software, 70(5). 10.18637/jss.v070.i05

Laman, E. A., Rooper, C. N., Turner, K., Rooney, S., Cooper, D. W., Zimmermann, M. 2018. Using species distribution models to describe essential fish habitat in Alaska. Canadian Journal of Fisheries and Aquatic Sciences, 75(8), 1230–1255. 10.1139/cjfas-2017-0181

Leising, A., Hunsicker, M. E., Tolimieri, N. T., Williams, G., Harley, A. 2024. 2023-2024 California Current Ecosystem Report. NOAA California Current Integrated Ecosystem Assessment Team, Pacific Fishery Management Council, 10.25923/vxen-pf76.

Lewison, R. L., Freeman, S. A., Crowder, L. B. 2004. Quantifying the effects of fisheries on threatened species: the impact of pelagic longlines on loggerhead and leatherback sea turtles. Ecology Letters, 7(3), 221–231. 10.1111/j.1461-0248.2004.00573.x

Lindgren, F., Rue, H. 2015. Bayesian spatial modelling with *R*-INLA. Journal of Statistical Software, 63(19). 10.18637/jss.v063.i19

Lindgren, F., Rue, H., Lindström, J. 2011. An explicit link between Gaussian fields and Gaussian Markov Random Fields: The stochastic partial differential equation approach. Journal of the Royal Statistical Society Series B: Statistical Methodology, 73(4), 423–498. 10.1111/j.1467-9868.2011.00777.x

Melo-Merino, S. M., Reyes-Bonilla, H., Lira-Noriega, A. 2020. Ecological niche models and species distribution models in marine environments: A literature review and spatial analysis of evidence. Ecological Modelling, 415, 108837. 10.1016/j.ecolmodel.2019.108837

Methot, R. D., Wetzel, C. R. 2013. Stock synthesis: A biological and statistical framework for fish stock assessment and fishery management. Fisheries Research, 142, 86–99. 10.1016/j.fishres.2012.10.012

Ono, K., Katara, I., Eliasen, S. K., Broms, C., Campbell, A., dos Santos Schmidt, T. C., Egan, A., Hølleland, S. N., Jacobsen, J. A., Jansen, T., Mackinson, S., Mousing, E. A., Nash, R. D. M., Nikolioudakis, N., Nnanatu, C., Nøttestad, L., Singh, W., Slotte, A., Wieland, K., Olafsdottir, A. H. 2024. Effect of environmental drivers on the spatiotemporal distribution of mackerel at age in the Nordic Seas during 2010-20. ICES Journal of Marine Science, 81(7), 1282–1294. 10.1093/icesjms/fsae087

Ohlberger, J., Langangen, Ø., Chr. Stige, L. 2022. Age structure affects population productivity in an exploited fish species. Ecological Applications 32(5), e2614. 10.1002/eap.2614

Olsen, E.M., Heupel, M.R., Simpfendorfer, C.A., Moland, E. 2012, Harvest selection on Atlantic cod behavioral traits: implications for spatial management. Ecology and Evolution, 2: 1549–1562. 10.1002/ece3.244

Ono, K., Shelton, A. O., Ward, E. J., Thorson, J. T., Feist, B. E., Hilborn, R. 2016. Space-time investigation of the effects of fishing on fish populations. Ecological Applications, 26(2), 392–406.

Phillips, N. D., Reid, N., Thys, T., Harrod, C., Payne, N. L., Morgan, C. A., White, H. J., Porter, S., Houghton, J. D. R. 2017. Applying species distribution modelling to a data poor, pelagic fish complex: the ocean sunfishes. Journal of Biogeography, 44(10), 2176–2187. 10.1111/jbi.13033

R Core Team. 2024. R: A language and environment for statistical computing. R Foundation for Statistical Computing. https://www.R-project.org/

Rijnsdorp, A. D., Bolam, S. G., Garcia, C., Hiddink, J. G., Hintzen, N. T., Denderen, P. D. van, Kooten, T. van. 2018. Estimating sensitivity of seabed habitats to disturbance by bottom trawling based on the longevity of benthic fauna. Ecological Applications, 28(5), 1302– 1312. 10.1002/eap.1731

Thorson, J. T., Shelton, A. O., Ward, E. J., Skaug, H. J. 2015. Geostatistical delta-generalized linear mixed models improve precision for estimated abundance indices for West Coast groundfishes. ICES Journal of Marine Science, 72(5), 1297–1310. 10.1093/icesjms/fsu243

Thorson, J. T. 2019. Guidance for decisions using the Vector Autoregressive Spatio-Temporal (VAST) package in stock, ecosystem, habitat and climate assessments, Fisheries Research, 210, 143–161. 10.1016/j.fishres.2018.10.013

Tolimieri, N., Haltuch, M. 2023. Sea-level index of recruitment variability improves assessment model performance for sablefish /textitAnoplopoma fimbria. Canadian Journal of Fisheries and Aquatic Sciences, 80(6), 1006–1016.

Tolimieri, N., Haltuch, M. A., Lee, Q., Jacox, M. G., Bograd, S. J. 2018. Oceanographic drivers of sablefish recruitment in the California Current. Fisheries Oceanography, 27(5), 458–474. 10.1111/fog.12266

Tsikliras AC, Polymeros K. 2014. Fish market prices drive overfishing of the ‘big ones’. PeerJ 2:e638 10.7717/peerj.638

Werner, E. E., Gilliam, J. F. 1984. The ontogenetic niche and species interactions in size-structured populations. Annual Review of Ecology and Systematics, 15(1), 393–425. 10.1146/annurev.es.15.110184.002141

