## Supplementary Material for "Advancing spatially explicit fisheries management with age-specific species distribution models"

Table S1: Distribution of samples by age (columns) and years (rows) of female sablefish collected and aged by the West Coast Groundfish Bottom Trawl Survey, 2003 – 2023.

Table S2: Distribution of samples by age (columns) and years (rows) of female Pacific hake collected and aged by the West Coast Groundfish Bottom Trawl Survey, 2003 – 2019.

Figure S1: Estimated spatial parameters (range, and spatial and spatiotemporal standard deviations). Points represent mean estimates; lines represent 95% confidence intervals.

Figure S2: Estimated spatial catch per unit effort (CPUE, kg per km2) for Pacific hake; rows represent ages (1-5). CPUE has been scaled for each cohort for visualization purposes.

Figure S3: Estimated spatial catch per unit effort (CPUE, kg per km2) for sablefish; rows represent ages (0-4). CPUE has been scaled for each cohort for visualization purposes.

Figure S4: Time-varying coefficients (with random effects in year) relating predicted densities of age a fish in year t to observed numbers the following year.

Table S1

| **Year** | **0** | **1** | **2** | **3** | **4** | **5** | **6** | **7** | **8** | **9** |
| --- | --- | --- | --- | --- | --- | --- | --- | --- | --- | --- |
| **2003** | 16 | 75 | 59 | 209 | 140 | 31 | 18 | 14 | 20 | 9 |
| **2004** | 63 | 32 | 30 | 55 | 130 | 74 | 33 | 8 | 7 | 9 |
| **2005** | 1 | 118 | 41 | 103 | 98 | 131 | 52 | 29 | 6 | 4 |
| **2006** | 1 | 6 | 39 | 31 | 65 | 78 | 111 | 71 | 26 | 13 |
| **2007** | 1 | 28 | 5 | 62 | 27 | 47 | 65 | 100 | 63 | 19 |
| **2008** | 8 | 3 | 26 | 4 | 19 | 16 | 42 | 62 | 89 | 36 |
| **2009** | 10 | 167 | 9 | 10 | 7 | 23 | 12 | 18 | 20 | 54 |
| **2010** | 12 | 11 | 201 | 22 | 9 | 7 | 19 | 18 | 21 | 24 |
| **2011** | 19 | 112 | 24 | 169 | 19 | 8 | 13 | 18 | 15 | 20 |
| **2012** | 20 | 47 | 78 | 43 | 113 | 27 | 10 | 5 | 7 | 7 |
| **2013** | 17 | 20 | 35 | 73 | 57 | 80 | 16 | 6 | 7 | 6 |
| **2014** | 14 | 177 | 15 | 42 | 50 | 50 | 56 | 24 | 14 | 5 |
| **2015** | 5 | 49 | 170 | 62 | 32 | 53 | 31 | 34 | 11 | 4 |
| **2016** | 20 | 14 | 52 | 160 | 48 | 41 | 41 | 37 | 23 | 18 |
| **2017** | 7 | 124 | 33 | 61 | 101 | 39 | 27 | 27 | 24 | 8 |
| **2018** | 11 | 26 | 168 | 46 | 67 | 99 | 49 | 50 | 38 | 22 |
| **2019** | 15 | 45 | 7 | 103 | 26 | 18 | 73 | 18 | 20 | 13 |
| **2020** | No survey | | | | | | | | | |
| **2021** | 61 | 241 | 72 | 34 | 22 | 169 | 57 | 47 | 41 | 34 |
| **2022** | 4 | 165 | 197 | 64 | 18 | 14 | 100 | 22 | 19 | 18 |
| **2023** | 55 | 20 | 185 | 193 | 42 | 29 | 34 | 89 | 19 | 17 |

Table S2

| **Year** | **1** | **2** | **3** | **4** | **5** |
| --- | --- | --- | --- | --- | --- |
| **2003** | 161 | 124 | 70 | 585 | 158 |
| **2004** | 0 | 0 | 0 | 0 | 0 |
| **2005** | 0 | 0 | 0 | 0 | 0 |
| **2006** | 0 | 0 | 0 | 0 | 0 |
| **2007** | 121 | 296 | 42 | 199 | 33 |
| **2008** | 89 | 200 | 216 | 44 | 152 |
| **2009** | 228 | 42 | 99 | 95 | 17 |
| **2010** | 79 | 99 | 25 | 63 | 105 |
| **2011** | 179 | 50 | 65 | 14 | 44 |
| **2012** | 47 | 170 | 40 | 52 | 18 |
| **2013** | 47 | 13 | 96 | 17 | 23 |
| **2014** | 70 | 53 | 15 | 148 | 30 |
| **2015** | 159 | 49 | 34 | 28 | 167 |
| **2016** | 38 | 139 | 46 | 40 | 38 |
| **2017** | 126 | 38 | 97 | 14 | 58 |
| **2018** | 103 | 75 | 15 | 116 | 14 |
| **2019** | 24 | 40 | 30 | 5 | 38 |

Figure S1

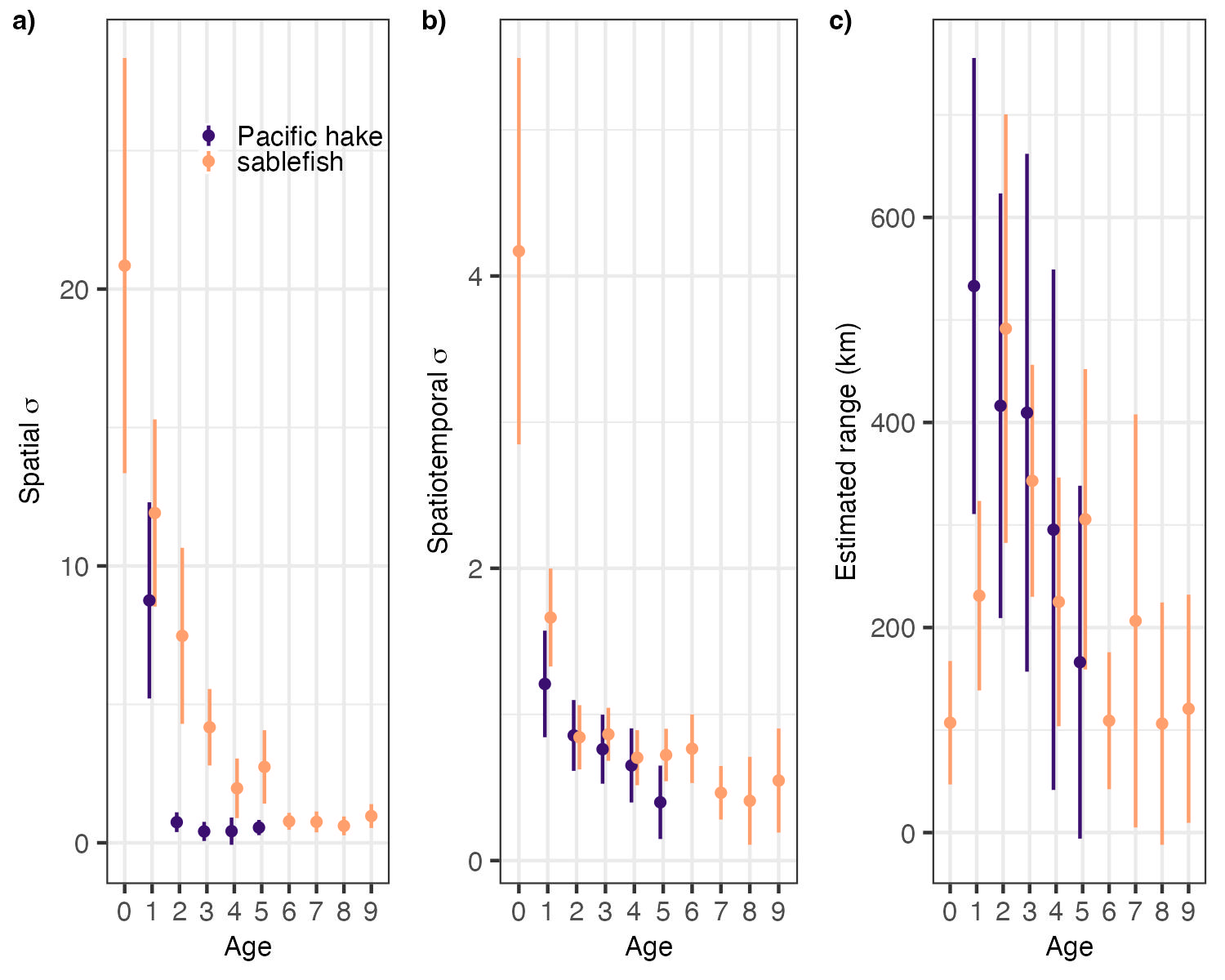

Figure S2

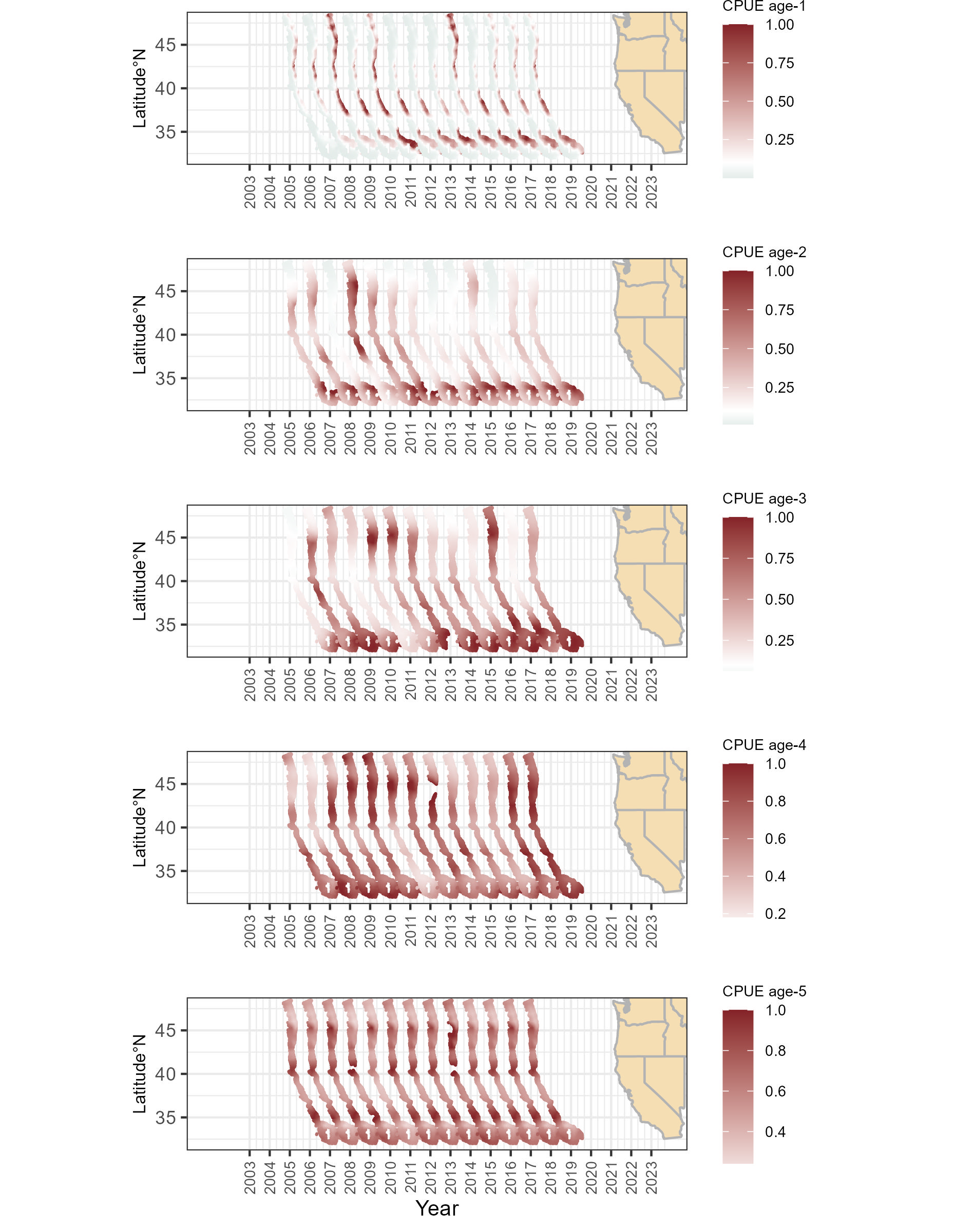

Figure S3

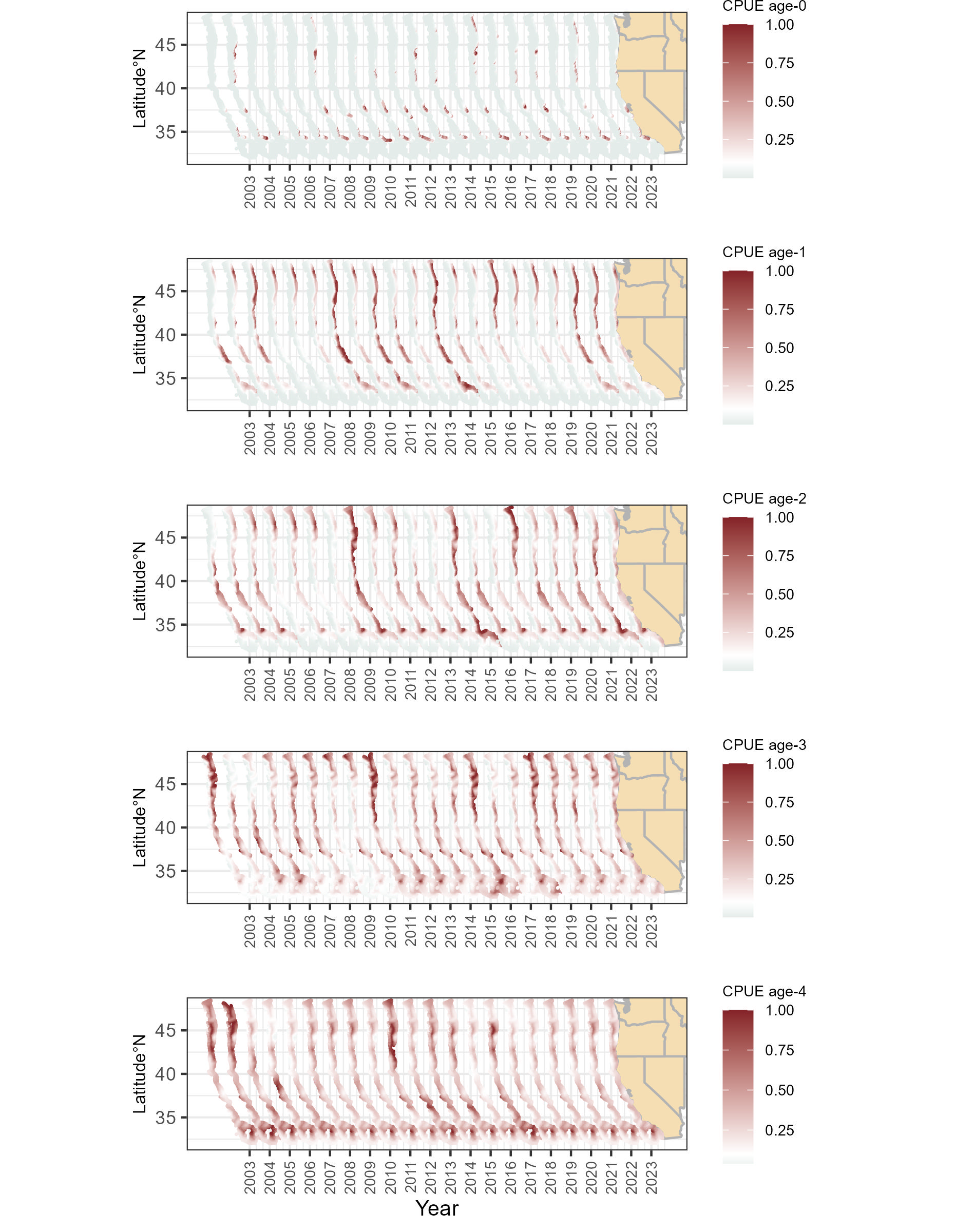

Figure S4

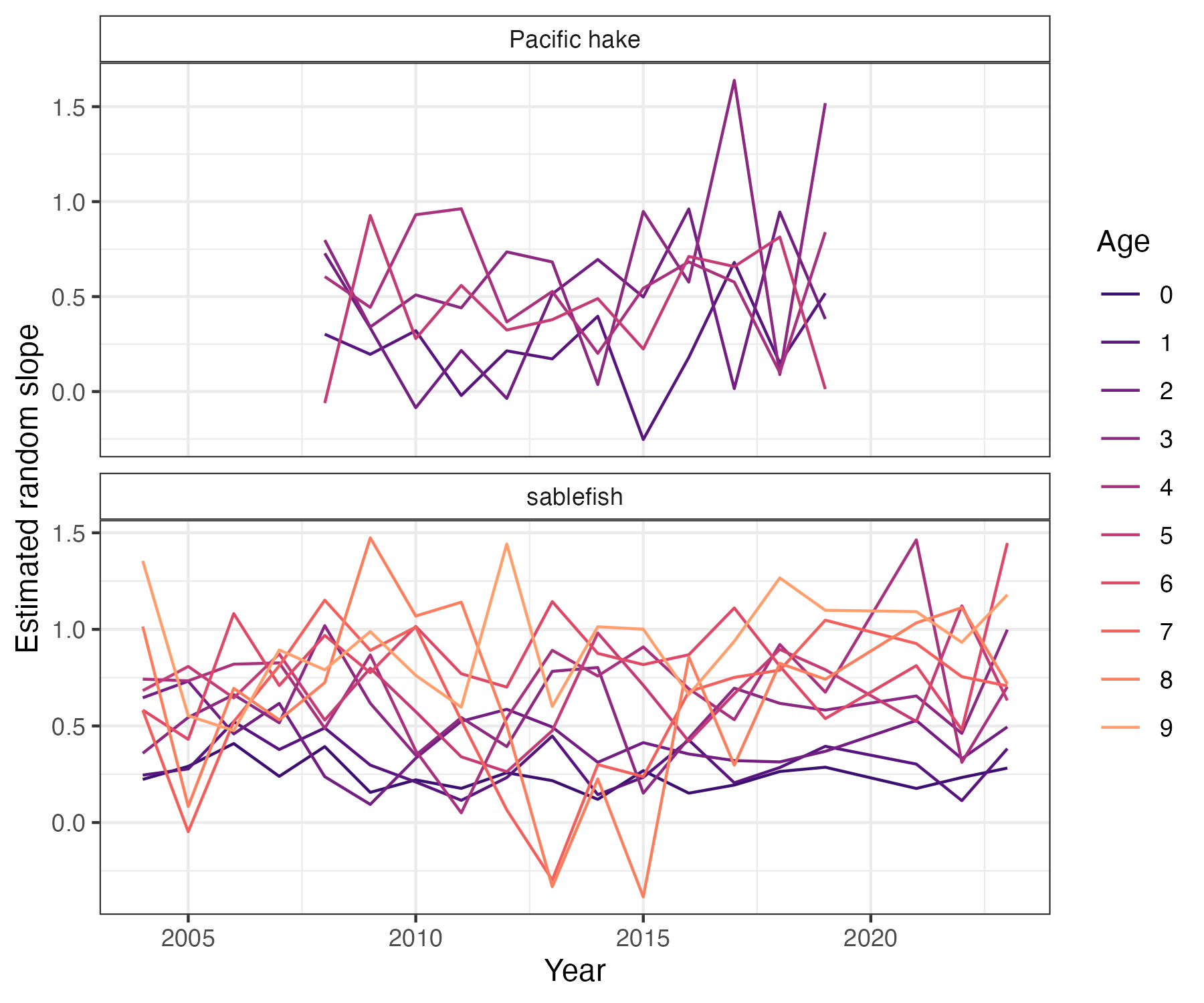
